# Neighborhood-Level Disadvantage Impacts Multiple Measures of Brain Health: An Imaging Epidemiology Study

**DOI:** 10.64898/2026.03.16.712147

**Authors:** Ethan H. Willbrand, Elizabeth M. Stoeckl, Daryn Belden, Sheena Y. Chu, Eleanna M. Melcher, Daniil Zhitnitskii, Elena Bonke, Jussi Mattila, Usman Iftikhar, Juha Koikkalainen, Antti Tolonen, Jyrki Lötjönen, Richard Bruce, John-Paul J. Yu

## Abstract

**Background:** The relationship between neighborhood-level socioeconomic disadvantage and brain health is an emerging area of research with critical implications for public health and clinical practice, yet its influence on brain structure remains unclear.

**Purpose:** To investigate the epidemiological association between neighborhood-level socioeconomic disadvantage [Area Deprivation Index (ADI)] and morphometric neuroimaging variables in a consecutive, non-disease enriched patient population.

**Materials and Methods:** This study, conducted at an academic medical center and associated community partners, used consecutive cross-sectional MRI neuroimaging data from 2,826 inpatient and outpatient individuals without radiological evidence of disease from January 2024 to June 2024. ADI, a geospatially determined index of neighborhood-level disadvantage, was calculated for each individual. Linear regressions tested the relationship between ADI and multiple morphometric variables: brain age gap (BAG; estimated - chronological BA), total brain tissue volume (TBV; total gray + white matter), five subcortical region volumes (hippocampus, thalamus, caudate, putamen, and nucleus accumbens) and four cortical region volumes [anterior cingulate cortex, posterior cingulate cortex, medial prefrontal cortex (MPFC), lateral PFC (LPFC)]. Volumetric measures were normalized to intracranial volume. Models controlled for age, sex, and total white matter hyperintensity volume (WMHV).

**Results:** 2,826 individuals (mean age, 52.7 ± 18.8 [standard deviation]; 1732 women) were evaluated. Residence in the 20% most disadvantaged neighborhoods was associated with a higher BAG (βs > 2.12, Ps < .01) and decreased TBV (βs < −5.12, Ps < .05). Additionally, increased WMHV was higher among those in the most disadvantaged neighborhoods (ts < - 2.50, Ps < .05) and associated with lower volume in most regions. Interaction models showed increased negative associations between WMHV and volumes of the caudate, nucleus accumbens, and lateral prefrontal cortex among those in the most disadvantaged neighborhoods.

**Conclusions:** Neighborhood disadvantage is associated with adverse brain morphometry, including higher BAG, lower TBV, and amplified vascular-related regional volume loss.

**Key Results:**

- In 2,826 adults (mean age, 53 years ± 19; 1,732 women), residence in the most disadvantaged neighborhoods (national: 116/2,826, 4%; state: 129/2826, 5%) was associated with higher brain age gap at the national (β = 2.12, 95% CI = 0.81 to 3.43, *P* = .001) and state levels (β = 2.36; 95% CI = 1.10 to 3.61, *P* < .001).
- Total brain tissue volume was lower at the national (β = −5.12, 95% CI = −10.13 to −0.11, *P* = .045) and state levels (β = −6.13, 95% CI = −10.90 to −1.37, *P* = .011).
- White matter hyperintensity volume was higher in the most disadvantaged group (national: *P* = .013; state: *P* = .003) and demonstrated amplified associations with caudate, nucleus accumbens, and lateral prefrontal cortex volumes in the most disadvantaged group at the national and/or state levels (*P*s < .05).

## Introduction

The relationship between neighborhood-level socioeconomic disadvantage and brain health is an emerging area of research with critical implications for basic research, public health, and clinical practice. The life-course social exposome—the conditions in which individuals are born, live, work, and age—profoundly shape long-term health outcomes (1,2). Within this framework, neighborhood disadvantage, quantified by indices such as the Area Deprivation Index (ADI), represents a particularly important determinant because it reflects accumulated structural inequities that extend beyond individual behaviors (2). These inequities manifest in limited access to quality healthcare, suboptimal nutrition, reduced opportunities for educational and occupational advancement, unsafe built environments, and heightened psychosocial stress (2). Over the life-course, such factors are increasingly recognized to contribute to disparities in clinical outcomes (3). Indeed, while these social exposome exposures have been linked to adverse outcomes ranging from neurocognitive development to cardio- and cerebrovascular burden to accelerated cognitive decline (3–5), their associations with morphometric brain markers in real-world clinical populations remain understudied (6–11).

To date, most neuroimaging studies examining socioeconomic disparities rely on enriched research cohorts or disease-specific samples, such as aging populations or children at neurodevelopmental risk (6–10). These investigations have revealed links between socioeconomic disadvantage and/or neighborhood-level deprivation and measures like smaller whole brain and regional volumes. However, the extent to which neighborhood disadvantage shapes brain structure in the context of everyday clinical care, across both inpatients and outpatients and outside of disease-enriched samples, has not been systematically investigated. Importantly, general clinical populations are inherently heterogeneous, spanning broad age ranges and diagnostic indications, and therefore provide a unique opportunity to assess whether neighborhood context exerts measurable neuroanatomical effects outside of curated cohorts.

Concomitantly, there is growing interest in brain age estimation models, which provide a biologically informed metric of overall brain health. Elevated brain age has been associated with cardiovascular risk factors, psychiatric illness, and cognitive impairment (12–14). Because brain age integrates distributed patterns of cortical and subcortical atrophy into a single biologically informed metric, it may be particularly sensitive to cumulative environmental and cardiometabolic exposures. However, whether neighborhood-level disadvantage is independently associated with increased brain age, and how it interacts with known imaging correlates of vascular pathology such as white matter hyperintensity volume (WMHV), remains an open question (15).

In this cross-sectional neuroimaging epidemiology study of 2,826 patients undergoing routine clinical magnetic resonance imaging (MRI), we tested whether living in the most socioeconomically disadvantaged neighborhoods was associated with three key classes of brain health metrics: i) higher brain age gap (BAG), ii) decreased total brain tissue volume (TBV), and iii) lower volumes across cortical and subcortical regions of interest implicated in aging and neuropsychiatric disease. We also explored whether WMHV [a neuroradiologic marker of small vessel disease and hypertension (16–19)] interacted with neighborhood disadvantage to compound adverse neuroanatomical changes (20). By leveraging a large, demographically diverse clinical sample not enriched for any specific disease, this study aims to uncover how structural inequity becomes biologically embedded in the brain and to establish neighborhood context as a critical covariate in both clinical interpretation and population-level neuroimaging research.

## Materials and Methods

### Study Participants

Neuroimaging data were acquired from consecutive inpatient and outpatient clinical MRI examinations from January 2024 to June 2024. Originally, data from 35,704 patients were acquired (**Fig 1**). Patients were first excluded if their imaging impression indicated any characteristic of disease (e.g., stroke, tumor/mass, demyelination) or they were less than 18 years old (**Fig 1**). Next, patients were excluded if we were unable to acquire ADI or morphometric measures for them (described in subsequent sections; **Fig 1**). This resulted in a final sample of 2,826 patients (**Fig 1**).

**Figure 1:**
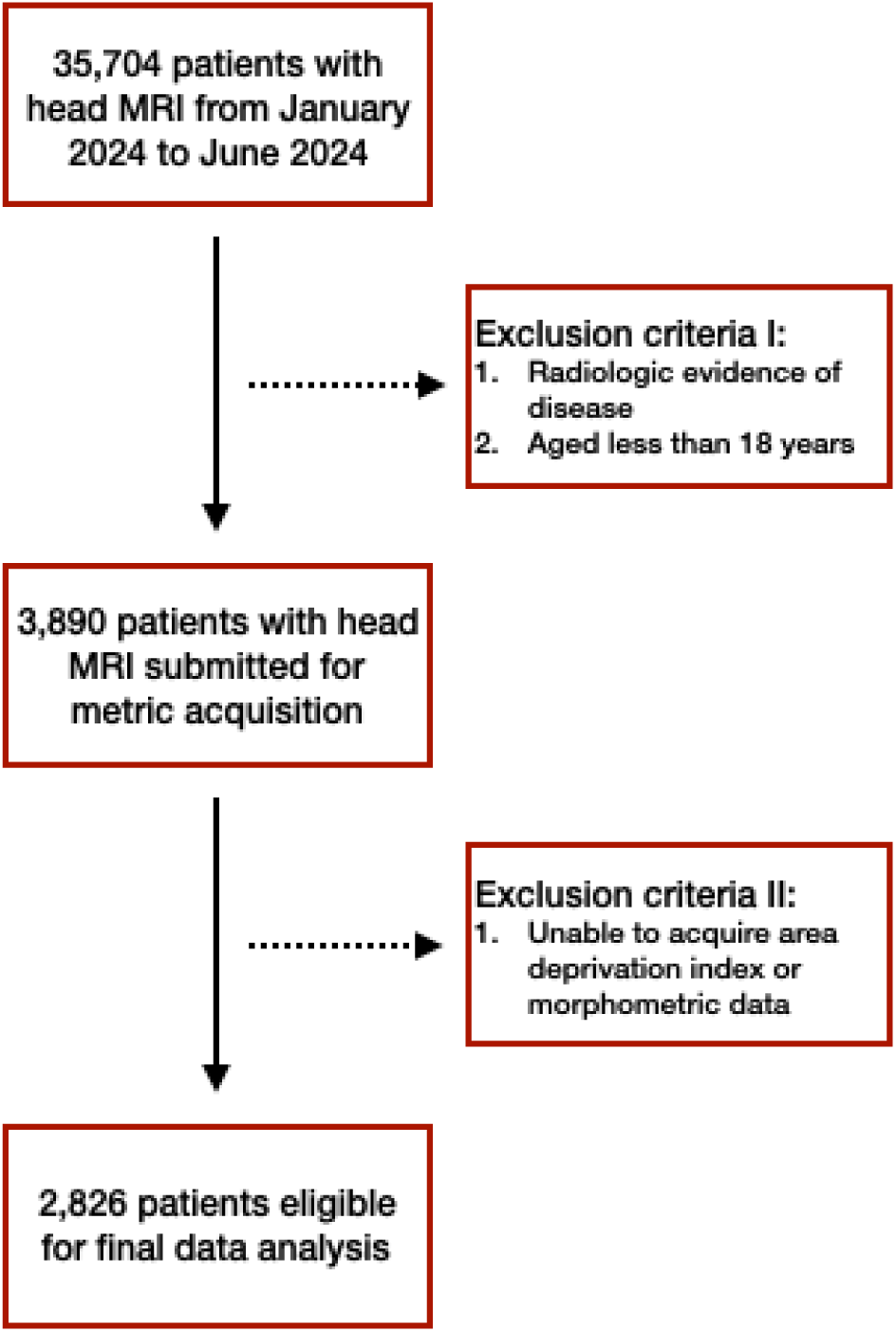
Flowchart outlining patient selection. Leftward boxes connected by solid arrows indicate the number of patients at each exclusion checkpoint. Rightward boxes stemming from dashed arrows highlight exclusion criteria.

### Neighborhood Disadvantage

Neighborhood disadvantage was quantified using the ADI, a census block group–level composite measure incorporating 17 indicators of income, education, employment, and housing characteristics derived from 2023 American Community Survey data (2). Detailed methodology and component variables have been described previously (21), and ADI scores are publicly available via the Neighborhood Atlas (2). Census block groups are assigned state- and national-level rankings to allow relative comparison of disadvantage. Participants’ most recent residential addresses were geocoded using ZIP+4 identifiers and linked to corresponding census block groups to assign ADI rankings (**Fig 2**).

**Figure 2:**
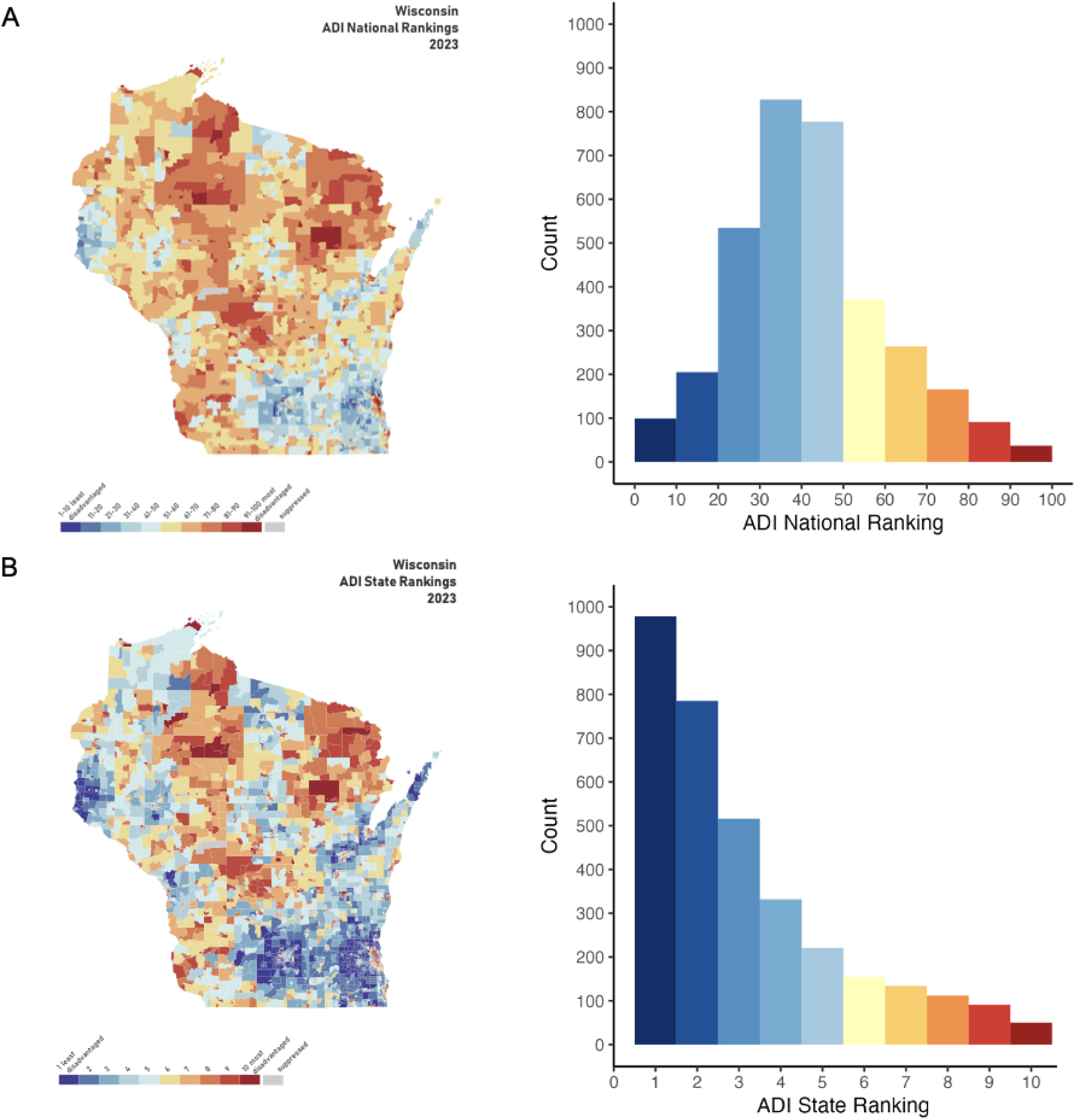
National and state neighborhood socioeconomic disadvantage rankings of the study patients. **(A)** Left: Area deprivation scores mapped onto the state of Wisconsin, where the study sample resides, with national ranking deciles shown (from: https://www.neighborhoodatlas.medicine.wisc.edu/). Right: Histogram quantifying the counts of participants in each area deprivation index decile. **(B)** Same as A, but for state ranking deciles. Abbreviations: area disadvantage index (ADI).

### Morphometric Data Acquisition

Structural brain morphometry was derived from T1-weighted MRIs from each participant using an automated, atlas-based segmentation pipeline (cNeuro, cMRI, Combinostics Oy, Tampere, Finland). T1 images were segmented into cortical and subcortical regions using a validated multi-atlas framework in which subject images are nonlinearly registered to a reference atlas set (https://www.neuromorphometrics.com/) and regional labels are propagated and fused to generate final segmentations. This approach has been previously validated for extraction of hippocampal and whole-brain volumes in neurodegenerative disease populations (22–24). Regional volumes were computed as the voxel count within each labeled region multiplied by voxel dimensions. Composite volumetric measures were derived from the corresponding regional segmentations. To account for interindividual differences in head size, volumetric measures were normalized using an intracranial volume–based scaling approach (25). Age- and sex-adjusted percentiles were generated using reference distributions derived from large normative cohorts, as previously described (25). WMHV was segmented from T2-weighted FLAIR images using an automated method adapted from previously validated structural MRI approaches (23). Estimated brain age was calculated using a convolutional neural network leveraging probabilistic tissue segmentations defined from T1 images. The model was trained using T1 images from 7,578 subjects.

### Study Variables

Based on prior work (7), a dichotomous ADI variable was created using the highest quintile (most disadvantaged neighborhoods) vs the lowest four quintiles (least disadvantaged neighborhoods) based on the relative national and state ADI distributions. We tested the following neuroimaging metrics given their associations with aging and numerous neurological and psychiatric disorders: BAG (estimated BA - actual BA), TBV (sum of total gray and white matter), five subcortical regions (hippocampus, thalamus, caudate, putamen, and nucleus accumbens) and four cortical regions [anterior cingulate cortex (ACC), posterior CC (PCC), medial prefrontal cortex (MPFC), lateral PFC (LPFC)]. Sex and age were included as covariates. Main effects and interactions with WMHV were specifically assessed given that this measure is a well-characterized proxy for small vessel cerebrovascular disease and hypertension (16–19). Sex was included as a dichotomous variable in all analyses, morphometric measures and age were treated as continuous variables.

### Statistical Analysis

All statistical analyses were conducted using R statistical software version 4.1.2. Given the large sample size, the D’Agostino skewness test was used to assess the distribution of both ADI metrics. Categorical demographic control variables were tested against ADI with Pearson’s Chi-squared (χ²) test. Continuous control variables were tested against ADI with Welch two sample t-tests. Linear regressions were fit to test associations between the morphometric brain metrics and national ADI, including sex, age, and WMHV as covariates. We used the standard significance level for hypothesis testing (two-sided α = .05) in all statistical tests. All analyses were also conducted with the state ADI distributions to assess whether the scale of index impacted the results. We also conducted sensitivity analyses by treating ADI as a continuous variable. All models that showed a significant association between ADI and morphometric measures were internally validated by bootstrapping with 5,000 resamples. Finally, we implemented exploratory models to test for interactions between our three covariates and ADI on our morphometric measures.

## Results

### Demographic Characteristics

Full demographic characteristics of the patients broken down by ADI level are detailed in **Table 1** (National ADI) and **Table S1** (State ADI). The ages and ethnicities of patients living in the most disadvantaged neighborhoods did not significantly differ from those living in the least disadvantaged neighborhoods (all *P*s > .05); however, sex did (National: χ²(1) = 4.25, *P* = .039; **Table 1**; State: χ²(1) = 7.26, *P* = .007; **Table S1**). Notably, the distributions of both the national-and state-level ADI rankings of the patients were right-skewed (National: Skew = 0.55, Z = 12.23, *P* < .001; State: Skew = 1.19, Z = 22.79, *P* < .001; **Fig 2**), indicating that the hospital in this sample was primarily serving participants in lower disadvantaged neighborhoods.

**Table 1:**
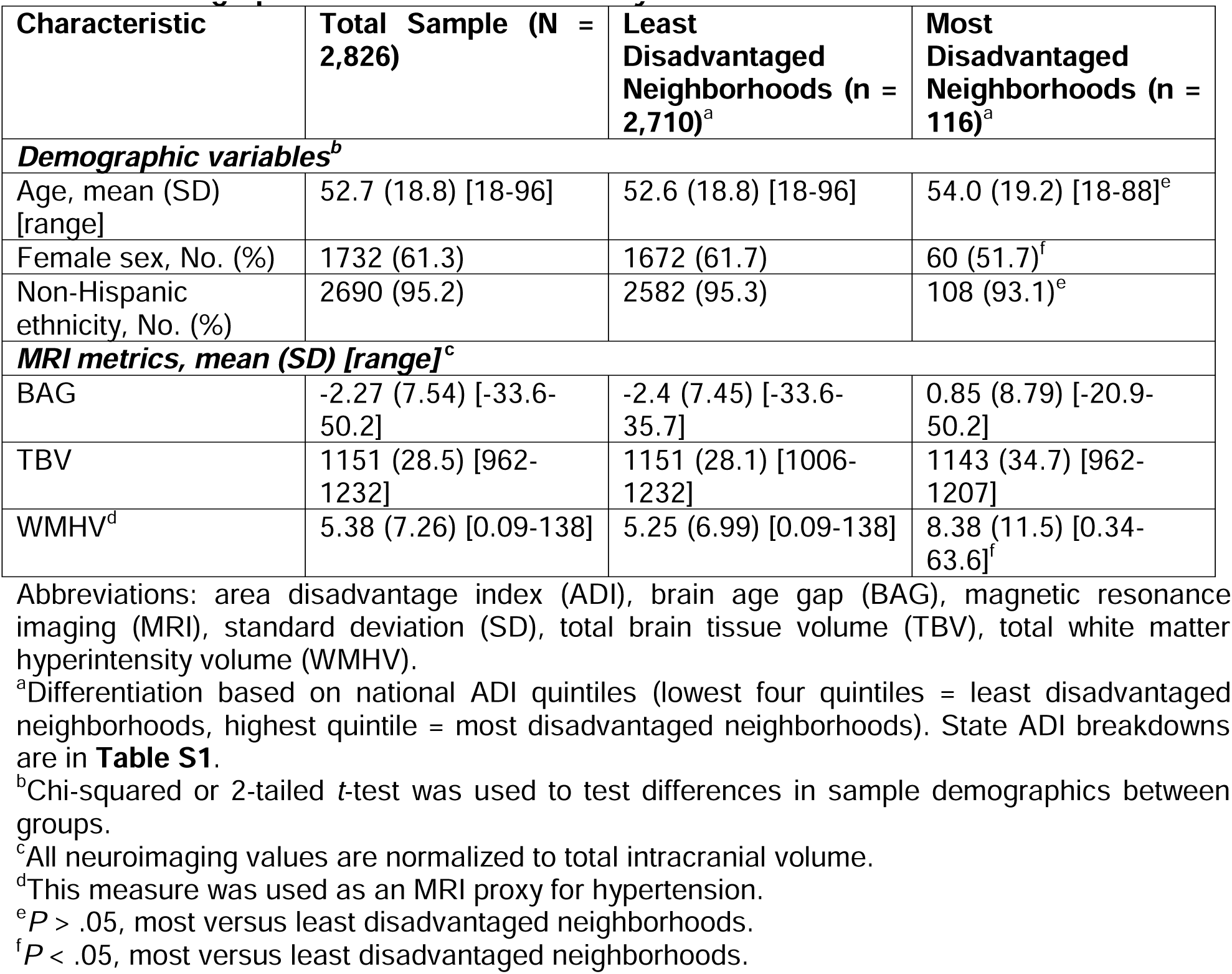
Demographic Characteristics of Study Patients.

### Neighborhood-Level Disadvantage and Brain Age Gap

We first tested for associations between neighborhood-level disadvantage and BAG in our patient sample. Linear regression analysis identified that living in the most disadvantaged neighborhoods at the national (β = 2.12, 95% CI = 0.81 to 3.43, *P* = .001; **Fig 3A**) and state (β = 2.36, 95% CI = 1.10 to 3.61, *P* < .001; **Fig S1A**) levels was associated with a higher BAG (**Table 2**). These results held for sensitivity analyses where neighborhood-level disadvantage was treated as a continuous variable instead of a dichotomous variable (**Table S2**). These models also showed that male sex and higher WMHV were also associated with a higher BAG (**Table 2**). However, neither of these variables interacted with ADI (state or national) to impact brain age (all *P*s > .05).

**Figure 3:**
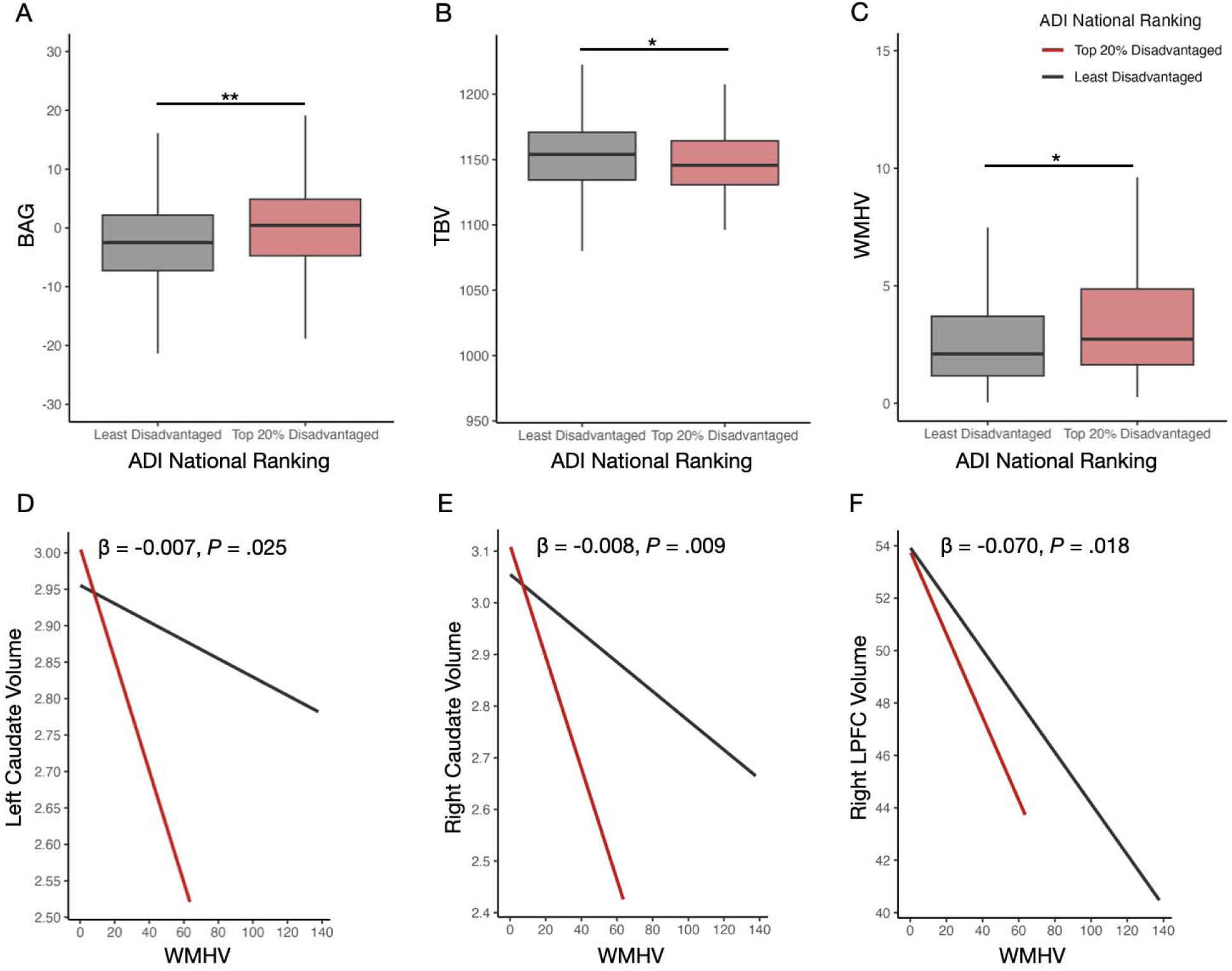
National neighborhood socioeconomic disadvantage is associated with multiple imaging measures of brain health. **(A)** Brain age gap (y-axis) as a function of area deprivation index national ranking (x-axis) as a boxplot. **(B)** Same as A, but for total brain tissue volume. **(C)** Same as B, but for total white matter hyperintensity volume. **(D)** Left caudate volume (y-axis) as a function of total white matter hyperintensity volume (x-axis) and ADI national ranking (color; see key). Best fit line is shown, along with the β and P-value for the interaction effect. **(E)** Same as D, but for right caudate volume. **(F)** Same as D, but for right lateral prefrontal cortex volume. All morphometric variables are normalized to intracranial volume. Asterisks indicate the following *P*-value thresholds: ** *P* < .01, * *P* < 0.05. Abbreviations: area disadvantage index (ADI), brain age gap (BAG), lateral prefrontal cortex (LPFC), total brain tissue volume (TBV), total white matter hyperintensity volume (WMHV).

**Table 2:** Regression Model Results for Association Between Neighborhood-Level Disadvantage and Brain Age Gap.

| Variable | Adjusted Model<sup>a</sup> |  | Bootstrapped Model<sup>b</sup> |  |
| --- | --- | --- | --- | --- |
| *National ADI* |  |  |  |  |
|  | β-value (95% CI)<sup>c</sup> | *P*-value | β-value (95% CI)<sup>c</sup> | *P*-value |
| 20% Most disadvantaged neighborhoods | 2.12 (0.81 to 3.43) | .0014 | 2.10 (0.77 to 3.51) | .027 |
| Male sex | 2.73 (2.20 to 3.26) | < .001 | 2.73 (2.20 to 3.26) | < .001 |
| WMHV | 0.29 (0.25 to 0.32) | < .001 | 0.29 (0.21 to 0.38) | < .001 |
| *State ADI* |  |  |  |  |
| 20% Most disadvantaged neighborhoods | 2.36 (1.10 to 3.61) | < .001 | 2.34 (1.02 to 3.68) | .011 |
| Male sex | 2.71 (2.18 to 3.24) | < .001 | 2.70 (2.19 to 3.24) | < .001 |
| WMHV | 0.29 (0.25 to 0.32) | < .001 | 0.29 (0.21 to 0.38) | < .001 |

Abbreviations: area disadvantage index (ADI), total white matter hyperintensity volume (WMHV).
<sup>a</sup>National ADI model: $F(3,2916) = 131.0$ , $P < .001$ , Adjusted $R^2 = 0.12$ ; State ADI model: $F(3,2916) = 132.3$ , $P < .001$ , Adjusted $R^2 = 0.12$ .
<sup>b</sup>National ADI model: Bootstrapped $R^2$ (95% CI) = 0.12 (0.09-0.16); State ADI model: Bootstrapped $R^2$ (95% CI) = 0.12 (0.09-0.16).
<sup>c</sup>Values presented are unstandardized.

### Neighborhood-Level Disadvantage and Whole-Brain-Level Morphometry

We next tested whether neighborhood-level disadvantage was associated with TBV. Here, linear regression analysis identified that living in the most disadvantaged neighborhoods at the national (β = −5.12, 95% CI = −10.13 to −0.11, *P* = .045; **Fig 3B**) and state (β = −6.13, 95% CI = - 10.90 to −1.37, *P* = .011; **Fig S1B**) levels was associated with a decrease in TBV (**Table 3**).

**Table 3:** Regression Model Results for Association Between Neighborhood-Level Disadvantage and Total Brain Tissue Volume.

| Variable | Adjusted Model<sup>a</sup> |  | Bootstrapped Model<sup>b</sup> |  |
| --- | --- | --- | --- | --- |
| ***National ADI*** |  |  |  |  |
|  | β-value (95% CI)<sup>c</sup> | *P*-value | β-value (95% CI)<sup>c</sup> | *P*-value |
| 20% Most disadvantaged neighborhoods | -5.12 (-10.13 to -0.11) | .045 | -5.10 (-11.57 to 0.77) | .17 |
| Male sex | -5.80 (-7.85 to -3.76) | < .001 | -5.80 (-7.91 to -3.68) | < .001 |
| Age | -0.42 (-0.47 to -0.36) | < .001 | -0.42 (-0.47 to -0.36) | < .001 |
| WMHV | -0.42 (-0.56 to -0.28) | < .001 | -0.43 (-0.72 to -0.17) | .0029 |
| ***State ADI*** |  |  |  |  |
| 20% Most disadvantaged neighborhoods | -6.13 (-10.90 to -1.37) | .011 | -6.08 (-12.35 to -0.50) | .099 |
| Male sex | -5.75 (-7.79 to -3.71) | < .001 | -5.75 (-7.85 to -3.62) | < .001 |
| Age | -0.42 (-0.47 to -0.36) | < .001 | -0.42 (-0.47 to -0.36) | < .001 |
| WMHV | -0.42 (-0.55 to -0.28) | < .001 | -0.43 (-0.72 to -0.17) | .0031 |

Abbreviations: area disadvantage index (ADI), total white matter hyperintensity volume (WMHV).
<sup>a</sup>National ADI model: $F(4,2822) = 88.32$ , $P < .001$ , Adjusted $R^2 = 0.11$ ; State ADI model: $F(4,2822) = 88.98$ , Adjusted $R^2 = 0.11$ .
<sup>b</sup>National ADI model: Bootstrapped $R^2$ (95% CI) = 0.11 (0.09-0.14); State ADI model: Bootstrapped $R^2$ (95% CI) = 0.11 (0.09-0.14).
<sup>c</sup>Values presented are unstandardized.

These results held for sensitivity analyses where neighborhood-level disadvantage was treated as a continuous variable instead of a dichotomous variable (**Table S3**). These models also showed that male sex, higher age, and higher WMHV were also associated with decreased TBV (**Table 3**). None of these variables interacted with ADI (state or national) to impact TBV (all *P*s > .05).

### Neighborhood-Level Disadvantage and ROI-Level Brain Morphometry

Finally, we examined the relationship between neighborhood-level disadvantage and the volumes of five subcortical regions (hippocampus, thalamus caudate, putamen, and nucleus accumbens) and four cortical regions (ACC, PCC, MPFC, and LPFC). ADI was not associated with any of these regions (**Table S4–12**). There were associations between male sex, age, and WMHV with the majority of these regional volumes (**Table S4–12**). Of note, WMHV was significantly increased in those living in the most disadvantaged neighborhoods (National: t(117.45) = −2.50, *P* = .013; **Fig 3C**; **Table 1**; State: t(132.71) = −2.96, *P* = .003; **Fig S1C**; **Table S1**). Given this, subsequent interaction models identified significant national ADI x WMHV interaction on left caudate volume (**Fig 3D**), right caudate volume (**Fig 3E**), and right LPFC volume (**Fig 3F**). These interactions were present for state ADI, with the additional ADI x WMHV interaction on left nucleus accumbens, and right nucleus accumbens, and left LPFC volumes (**Fig S1D–I**). There were no other significant interactions for any other region or with sex or age (all *P*s > .05).

## Discussion

### Associations between Neighborhood-Level Disadvantage and Brain Morphometry in a large, non-enriched research cohort

In this large cross-sectional neuroimaging study of over 2,800 inpatient and outpatient individuals without radiological evidence of disease, we found that neighborhood-level socioeconomic disadvantage, measured using the ADI, was associated with multiple measures of adverse brain health. Specifically, individuals living in the most disadvantaged neighborhoods exhibited significantly elevated BAG and reduced TBV. While no main effects of ADI were observed on specific cortical and subcortical regions, exploratory interaction analyses revealed that ADI exacerbated the deleterious effects of WMHV on volumes of the caudate, nucleus accumbens, and LPFC. Crucially, these effects were observed in a consecutive, real-world clinical sample rather than a disease-enriched research cohort, suggesting that the neuroanatomical correlates of structural inequity are detectable even outside traditional epidemiologic or longitudinal aging studies. This expands prior work (7,8,20) by demonstrating that neighborhood context is not merely a background sociodemographic variable but a measurable correlate of brain structure that is observed in routine clinical care. Together, these findings provide new evidence that neighborhood disadvantage independently and interactively affects brain morphometry in clinical populations.

Our results immediately extend the growing literature linking social determinants of health to neurobiological outcomes. Prior studies have shown that living in socioeconomically deprived neighborhoods is associated with increased cardiovascular disease burden, psychiatric morbidity, and accelerated cognitive decline (3–5). Extending this work, we demonstrate that such disadvantage is also associated with a higher BAG, which is a biomarker linked to multiple adverse outcomes including dementia, psychiatric disorders, and mortality (12–14). Further, these findings directly support prior work suggesting a potential relationship between BAG and ADI (15), by examining this relationship in the largest sample to date and, importantly, within a clinically unenriched real-world population, where we identify a robust association between higher ADI and higher BAG. The magnitude of brain age increase observed in our study empirically aligns with previous work suggesting that brain age reflects cumulative exposure to physiological and psychosocial stressors (26). Brain age models are increasingly conceptualized as integrative markers of multisystem biological wear-and-tear, capturing vascular, metabolic, inflammatory, and stress-related processes that converge on neuroanatomical decline (27–29). Thus, the association between ADI and elevated brain age may reflect the accumulation of chronic environmental adversity, including reduced access to healthcare, environmental toxins, psychosocial stress, and cardiometabolic risk.

We also found that neighborhood disadvantage was associated with reduced TBV, a finding consistent with prior studies in enriched research cohorts or disease-specific samples reporting lower gray and white matter volumes in socioeconomically disadvantaged children and adults (6–11,30). Most notably, these associations persisted in our sample after controlling for age, sex, and WMHV, and were robust across both national and state-level ADI indices. Because TBV reflects global cortical and subcortical integrity, these findings suggest that socioeconomic disadvantage on its own may influence diffuse neuroanatomical systems rather than regionally isolated processes. Such diffuse effects are consistent with models of chronic stress exposure, glucocorticoid dysregulation, and low-grade systemic inflammation, all of which have been implicated in accelerated brain aging and cortical thinning (31–33). Altogether, this emphasizes the potential for structural inequities to manifest as measurable changes in brain anatomy, even in real-world clinical populations not selected for or enriched in specific diseases.

### Neighborhood-Level Risk Augments the Association Between Vascular Burden and Regional Brain Morphometry

Although ADI alone did not significantly predict regional volumes in our primary models, which have been observed in the prior studies discussed above, the observed interactions between ADI and WMHV suggest a compounding effect of socioeconomic and vascular risk on subcortical and prefrontal regions. The interaction pattern observed here is especially poignant: WMHV was associated with lower regional volumes across the cohort, but this association was amplified among individuals living in the most disadvantaged neighborhoods. These brain areas, particularly the caudate and LPFC, are vulnerable to both vascular insults and chronic stress, and are implicated in executive function, reward processing, and mood regulation (31,34). This suggests that neighborhood-level adversity may potentiate the neuroanatomical consequences of vascular pathology, consistent with a “double-hit” model in which structural inequity and cardiometabolic risk synergistically accelerate neural vulnerability.

Mechanistically, hypertensive small vessel disease disproportionately affects frontostriatal circuits supplied by penetrating arterioles, which include the caudate and LPFC (16–18). Chronic psychosocial stress, in turn, has been shown to alter dendritic architecture and synaptic density within these same regions (31,34). The convergence of vascular compromise and stress-related neuroplastic changes may therefore explain the amplified volume reductions observed in disadvantaged neighborhoods. This interaction framework also helps reconcile why ADI did not demonstrate strong independent regional effects in primary models. Here, socioeconomic disadvantage may exert its most measurable neuroanatomical impact when coupled with vascular burden, rather than as a purely additive determinant. Altogether, these findings align with prior evidence that socioeconomic adversity may interact with cardiometabolic risk to influence brain structure and function (20,32,35).

### Limitations, Future Directions, and Clinical Implications

There are several limitations to discuss First, its cross-sectional nature limits causal inference. While the associations observed are robust, longitudinal data are needed to establish temporal relationships between neighborhood disadvantage and neuroanatomical change. Second, although we controlled for key covariates, unmeasured confounders, such as diet, physical activity, or educational attainment, may also contribute to the observed effects. Third, our cohort was drawn from a single academic medical center in Wisconsin that does not fully represent the broader national population. Finally, although WMHV was used as a proxy for vascular risk, we did not have direct measures of blood pressure, antihypertensive treatment, or cardiometabolic biomarkers. Future studies integrating imaging with clinical cardiovascular data will be critical to disentangle direct vascular effects from broader environmental pathways.

Despite these limitations, our study has important implications. It highlights the necessity of considering the social exposome, particularly neighborhood-level disadvantages, in neuroimaging research and clinical interpretation. Our findings suggest that morphometric measures, including brain age and TBV, may serve as intermediate biomarkers of structural inequity, with relevance for both population neuroscience and clinical risk stratification. From a translational perspective, incorporating neighborhood context into neuroimaging analyses may improve risk prediction models and help avoid misattributing environmentally driven variation to intrinsic neurodegenerative processes. Further, the interactive effects between ADI and WMHV point to potential synergies between socioeconomic and vascular pathways in brain aging, which may inform public health interventions. Future work should incorporate longitudinal imaging to track the effects of neighborhood disadvantage over time, integrate individual-level socioeconomic data, and explore mechanistic pathways (e.g., stress physiology, inflammation) linking environment to brain health. Ultimately, this line of investigation supports a biologically grounded model of health equity in which structural disadvantage becomes embedded in brain morphology, reinforcing the need for multilevel interventions spanning clinical care, community investment, and public health policy.

## Conclusions

In conclusion, our findings demonstrate that neighborhood socioeconomic disadvantage is associated with accelerated brain aging, reduced brain tissue volume, and increased susceptibility to cerebrovascular burden in key brain regions. These results highlight the role of structural inequities as biologically embedded determinants of brain health and advocate for the inclusion of neighborhood context in both neuroimaging research and clinical practice.

## Summary Statement

Residence in the most socioeconomically disadvantaged neighborhoods was associated with higher brain age gap, lower total brain volume, and vascular-related volume loss on MRI in a consecutive non-enriched clinical cohort.

## Abbreviations

ADI: Area Deprivation Index
ACC: anterior cingulate cortex
BAG: Brain age gap
LPFC: lateral prefrontal cortex
MPFC: medial prefrontal cortex
PCC: posterior cingulate cortex
TBV: Total brain tissue volume
WMHV: White matter hyperintensity volume

