## Supplemental Material for "Neighborhood-Level Disadvantage Impacts Multiple Measures of Brain Health: An Imaging Epidemiology Study"

**Figure S1:** State Neighborhood Socioeconomic Disadvantage is Associated with Multiple Imaging Measures of Brain Health.

**Table S1:** Demographic Characteristics of Study Patients

**Table S2:** Sensitivity Analysis: Regression Model Results for Association Between Neighborhood-Level Disadvantage as a Continuous Variable and Brain Age Gap

**Table S3:** Sensitivity Analysis: Regression Model Results for Association Between Neighborhood-Level Disadvantage as a Continuous Variable and Total Brain Tissue Volume

**Table S4:** Regression Model Results for Association Between Neighborhood-Level Disadvantage and Hippocampus Volume

**Table S5:** Regression Model Results for Association Between Neighborhood-Level Disadvantage and Thalamus Volume

**Table S6:** Regression Model Results for Association Between Neighborhood-Level Disadvantage and Caudate Volume

**Table S7:** Regression Model Results for Association Between Neighborhood-Level Disadvantage and Putamen Volume

**Table S8:** Regression Model Results for Association Between Neighborhood-Level Disadvantage and Nucleus Accumbens Volume

**Table S9:** Regression Model Results for Association Between Neighborhood-Level Disadvantage and Anterior Cingulate Cortex Volume

**Table S10:** Regression Model Results for Association Between Neighborhood-Level Disadvantage and Posterior Cingulate Cortex Volume

**Table S11:** Regression Model Results for Association Between Neighborhood-Level Disadvantage and Medial Prefrontal Cortex Volume

**Table S12:** Regression Model Results for Association Between Neighborhood-Level Disadvantage and Lateral Prefrontal Cortex Volume

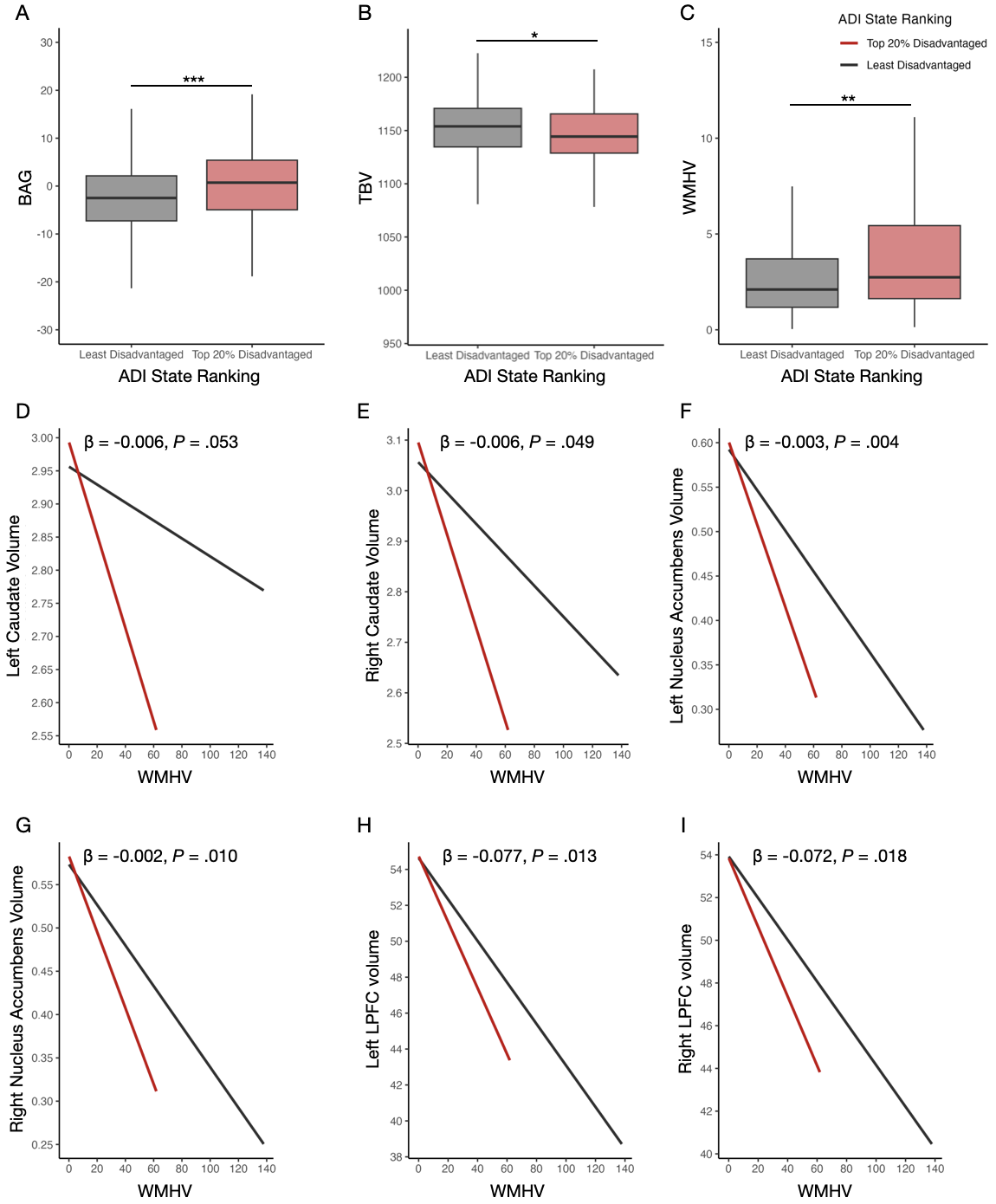

**Figure S1: State neighborhood socioeconomic disadvantage is associated with multiple imaging measures of brain health. (A)** Brain age gap (y-axis) as a function of area deprivation index state ranking (x-axis) as a boxplot. **(B)** Same as A, but for total brain tissue volume. **(C)** Same as A, but for total white matter hyperintensity volume. **(D)** Left caudate volume (y-axis) as a function of total white matter hyperintensity volume (x-axis) and area deprivation index state ranking (color; see key). Best fit line is shown, along with the β and *P*-value for the interaction effect. **(E)** Same as D, but for right caudate volume. **(F)** Same as D, but for left nucleus accumbens volume. **(G)** Same as D, but for right nucleus accumbens volume. **(H)** Same as D, but for left lateral prefrontal cortex volume. **(I)** Same as D, but for right lateral prefrontal cortex volume. All morphometric variables are normalized to intracranial volume. Asterisks indicate the following *P-*value thresholds: *** *P* < 0.001, ** *P* < .01, * *P* < 0.05. Abbreviations: area disadvantage index (ADI), brain age gap (BAG), lateral prefrontal cortex (LPFC), total brain tissue volume (TBV), total white matter hyperintensity volume (WMHV).

**Table S1: Demographic Characteristics of Study Patients**

| **Characteristic** | **Total Sample (N = 2,826)** | **Least Disadvantaged Neighborhoods (n = 2,697)**^a^ | **Most Disadvantaged Neighborhoods (n = 129)**^a^ |
| --- | --- | --- | --- |
| ***Demographic variables^b^*** | | | |
| Age, mean (SD) [range] | 52.7 (18.8) [18-96] | 52.6 (18.8) [18-96] | 53.2 (19.2) [18-92]^e^ |
| Female sex, No. (%) | 1732 (61.3) | 1672 (61.7) | 60 (51.7)^f^ |
| Non-Hispanic ethnicity, No. (%) | 2690 (95.2) | 2582 (95.3) | 108 (93.1)^e^ |
| ***MRI metrics, mean (SD) [range]* ^c^** | | | |
| BAG | -2.27 (7.54) [-33.6-50.2] | -2.43 (7.44) [-33.6-35.7] | 1.11 (8.72) [-18.8-50.2] |
| TBV | 1151 (28.5) [962-1232] | 1151 (28.1) [1006-1232] | 1143 (35.0) [962-1207] |
| WMHV^d^ | 5.38 (7.26) [0.09-138] | 5.24 (7.04) [0.09-138] | 8.29 (10.5) [0.16-61.9]^f^ |

Abbreviations: area disadvantage index (ADI), brain age gap (BAG), magnetic resonance imaging (MRI), standard deviation (SD), total brain tissue volume (TBV), total white matter hyperintensity volume (WMHV).

^a^Differentiation based on state ADI quintiles (lowest four quintiles = least disadvantaged neighborhoods, highest quintile = most disadvantaged neighborhoods).

^b^Chi-squared or 2-tailed *t*-test was used to test differences in sample demographics between groups.

^c^All neuroimaging values are normalized to intracranial volume .

^d^This measure was used as an MRI proxy for hypertension.

^e^*P* > .05, most versus least disadvantaged neighborhoods.

^f^*P* < .01, most versus least disadvantaged neighborhoods.

**Table S2: Sensitivity Analysis: Regression Model Results for Association Between Neighborhood-Level Disadvantage as a Continuous Variable and Brain Age Gap**

| **Variable** | **Adjusted Model**^a^ | | **Bootstrapped Model**^b^ | |
| --- | --- | --- | --- | --- |
| ***National ADI*** | | | | |
|  | β-value (95% CI)^c^ | *P*-value | β-value (95% CI)^c^ | *P*-value |
| Neighborhood-level disadvantage (continuous) | 0.019 (0.004-0.033) | .0098 | 0.018 (0.005-0.033) | .066 |
| Male sex | 2.73 (2.20-3.26) | < .001 | 2.73 (2.21-3.26) | < .001 |
| WMHV | 0.29 (0.25-0.32) | < .001 | 0.29 (0.21-0.38) | < .001 |
| ***State ADI*** | | | | |
| Neighborhood-level disadvantage (continuous) | 0.16 (0.051-0.27) | .0042 | 0.16 (0.049-0.28) | .044 |
| Male sex | 2.72 (2.19-3.25) | < .001 | 2.72 (2.20-3.25) | < .001 |
| WMHV | 0.29 (0.25-0.32) | < .001 | 0.29 (0.21-0.38) | < .001 |

Abbreviations: area disadvantage index (ADI), total white matter hyperintensity volume (WMHV).

^a^National ADI model: F(3,2916) = 129.7, *P* < .001, Adjusted R^2^ = 0.12; State ADI model: F(3,2916) = 130.3, *P* < .001, Adjusted R^2^ = 0.12.

^b^National ADI model: Bootstrapped R^2^ (95% CI) = 0.12 (0.09-0.15); State ADI model: Bootstrapped R^2^ (95% CI) = 0.12 (0.09-0.15).

^c^Values presented are unstandardized.

**Table S3: Sensitivity Analysis: Regression Model Results for Association Between Neighborhood-Level Disadvantage as a Continuous Variable and Total Brain Tissue Volume**

| **Variable** | **Adjusted Model**^a^ | | **Bootstrapped Model**^b^ | |
| --- | --- | --- | --- | --- |
| ***National ADI*** | | | | |
|  | β-value (95% CI)^c^ | *P*-value | β-value (95% CI)^c^ | *P*-value |
| Neighborhood-level disadvantage (continuous) | -0.072 (-10.13 to -0.11) | .0089 | -0.072 (-0.13 to -0.018) | .063 |
| Male sex | -5.77 (-7.81 to -3.72) | < .001 | -5.76 (-7.87 to -3.65) | < .001 |
| Age | -0.42 (-0.47 to -0.37) | < .001 | -0.42 (-0.47 to -0.36) | < .001 |
| WMHV | -0.41 (-0.55 to -0.27) | < .001 | -0.42 (-0.72 to -0.16) | .0041 |
| ***State ADI*** | | | | |
| Neighborhood-level disadvantage (continuous) | -0.52 (-0.94 to -0.091) | .017 | -0.52 (-0.96 to -0.073) | .095 |
| Male sex | -5.76 (-7.81 to -3.72) | < .001 | 5.76 (-7.88 to -3.66) | < .001 |
| Age | -0.42 (-0.47 to -0.36) | < .001 | -0.42 (-0.47 to -0.36) | < .001 |
| WMHV | -0.41 (-0.55 to -0.27) | < .001 | -0.42 (-0.72 to -0.16) | .0031 |

Abbreviations: area disadvantage index (ADI), total white matter hyperintensity volume (WMHV).

^a^National ADI model: F(4,2822) = 89.12, *P* < .001, Adjusted R^2^ = 0.11; State ADI model: F(4,2822) = 88.78, *P* < .001, Adjusted R^2^ = 0.11.

^b^National ADI model: Bootstrapped R^2^ (95% CI) = 0.11 (0.09-0.14); State ADI model: Bootstrapped R^2^ (95% CI) = 0.11 (0.09-0.14).

^c^Values presented are unstandardized.

**Table S4: Regression Model Results for Association Between Neighborhood-Level Disadvantage and Hippocampus Volume**

| **Variable** | **Left Hemisphere**^a^ | | **Right Hemisphere**^b^ | |
| --- | --- | --- | --- | --- |
| ***National ADI*** | | | | |
|  | β-value (95% CI)^c^ | *P*-value | β-value (95% CI)^c^ | *P*-value |
| 20% Most disadvantaged neighborhoods | 0.033 (-0.041 to 0.108) | .385 | 0.049 (-0.022 to 0.120) | .174 |
| Male sex | -0.019 (-0.050 to 0.011) | .216 | 0.0007 (-0.028 to 0.030) | .963 |
| Age | -0.002 (-0.003 to -0.001) | < .001 | -0.0008 (-0.002 to -0.000004) | .048 |
| MWHV | -0.006 (-0.008 to -0.004) | < .001 | -0.006 (-0.008 to -0.004) | < .001 |
| ***State ADI*** | | | | |
|  | β-value (95% CI)^c^ | *P*-value | β-value (95% CI)^c^ | *P*-value |
| 20% Most disadvantaged neighborhoods | 0.002 (-0.069 to 0.073) | .964 | 0.021 (-0.047 to 0.088) | .544 |
| Male sex | -0.019 (-0.049 to 0.012) | .227 | 0.001 (-0.028 to 0.030) | .947 |
| Age | -0.002 (-0.003 to -0.001) | < .001 | -0.0007 (-0.002 to -0.0000007) | .049 |
| WMHV | -0.006 (-0.008 to -0.004) | < .001 | -0.006 (-0.008 to -0.004) | < .001 |

Abbreviations: area disadvantage index (ADI), total white matter hyperintensity volume (WMHV).

^a^National ADI model: F(4, 2822) = 19.32, *P <* .001, Adjusted R^2^ = 0.03; State ADI model: F(4,2822) = 19.13, *P* < .001, Adjusted R^2^ = 0.03.

^b^National ADI model: F(4,2822) = 11.85, *P* < .001, Adjusted R^2^ = 0.02; State ADI model: F(4,2822) = 11.47, *P* < .001, Adjusted R^2^ = .02.

^c^Values presented are unstandardized.

**Table S5: Regression Model Results for Association Between Neighborhood-Level Disadvantage and Thalamus Volume**

| **Variable** | **Left Hemisphere**^a^ | | **Right Hemisphere**^b^ | |
| --- | --- | --- | --- | --- |
| ***National ADI*** | | | | |
|  | β-value (95% CI)^c^ | *P*-value | β-value (95% CI)^c^ | *P*-value |
| 20% Most disadvantaged neighborhoods | -0.044 (-0.053 to 0.142) | .376 | -0.035 (-0.063 to 0.131) | .484 |
| Male sex | -0.079 (-0.119 to -0.039) | < .001 | -0.130 (-0.170 to -0.091) | < .001 |
| Age | 0.001 (0.0002 to -0.002) | .016 | -0.003 (-0.004 to -0.002) | < .001 |
| WMHV | -0.014 (-0.017 to -0.012) | < .001 | -0.013 (-0.016 to -0.011) | < .001 |
| ***State ADI*** | | | | |
|  | β-value (95% CI)^c^ | *P*-value | β-value (95% CI)^c^ | *P*-value |
| 20% Most disadvantaged neighborhoods | -0.019 (-0.074 to 0.112) | .689 | 0.012 (-0.081 to 0.104) | .807 |
| Male sex | -0.079 (-0.119 to -0.039) | < .001 | -0.130 (-0.170 to –0.090) | < .001 |
| Age | -0.001 (0.0002 to 0.002) | .015 | -0.003 (-0.004 to -0.002) | < .001 |
| WMHV | -0.014 (-0.017 to -0.012) | < .001 | -0.013 (-0.016 to -0.011) | < .001 |

Abbreviations: area disadvantage index (ADI), total white matter hyperintensity volume (WMHV).

^a^National ADI model: F(4,2822) = 32.36, *P* < .001, Adjusted R^2^ = 0.04; State ADI model: F(4,2822) = 32.2, *P* < .001, Adjusted R^2^ = 0.04.

^b^National ADI model: F(4,2822) = 46.25, *P* < .001, Adjusted R^2^ = 0.06; State ADI model: F(4,2822) = 46.13, *P* < .001, Adjusted R^2^ = 0.06.

^c^Values presented are unstandardized.

**Table S6: Regression Model Results for Association Between Neighborhood-Level Disadvantage and Caudate Volume**

| **Variable** | **Left Hemisphere**^a^ | | **Right Hemisphere**^b^ | |
| --- | --- | --- | --- | --- |
| ***National ADI*** | | | | |
|  | β-value (95% CI)^c^ | *P*-value | β-value (95% CI)^c^ | *P*-value |
| 20% Most disadvantaged neighborhoods | 0.006 (to) | .852 | -0.0006 (-0.068 to 0.067) | .985 |
| Male sex | -0.057 (-0.057 to 0.070) | < .001 | -0.068 (-0.095 to -0.040) | < .001 |
| Age | -0.001 (-0.083 to -0.032) | < .001 | -0.001 (-0.002 to -0.0007) | < .001 |
| WMHV | -0.001 (-0.003 to 0.0003) | 0.102 | -0.003 (-0.005 to -0.001) | .001 |
| ***State ADI*** | | | | |
|  | β-value (95% CI)^c^ | *P*-value | β-value (95% CI)^c^ | *P*-value |
| 20% Most disadvantaged neighborhoods | -0.002 (-0.062 to 0.059) | .959 | -0.002 (-0.066 to 0.063) | .961 |
| Male sex | -0.057 (-0.083 to -0.031) | < .001 | -0.068 (-0.095 to -0.040) | < .001 |
| Age | -0.001 (-0.002 to -0.0005) | < .001 | -0.001 (-0.002 to -0.0007) | < .001 |
| WMHV | -0.001 (-0.003 to 0.0003) | 0.106 | -0.003 (-0.005 to -0.001) | .001 |

Abbreviations: area disadvantage index (ADI), total white matter hyperintensity volume (WMHV).

^a^National ADI model: F(4,2822) = 9.47, *P* < .001, Adjusted R^2^ = 0.01; State ADI model: F(4,2822) = 9.46, *P* < .001, Adjusted R^2^ = 0.01.

^b^National ADI model: F(4,2822) = 14.06, *P* < .001, Adjusted R^2^ = 0.02; State ADI model: F(4,2822) = 14.06, *P* < .001, Adjusted R^2^ = 0.02.

^c^Values presented are unstandardized.

**Table S7: Regression Model Results for Association Between Neighborhood-Level Disadvantage and Putamen Volume**

| **Variable** | **Left Hemisphere**^a^ | | **Right Hemisphere**^b^ | |
| --- | --- | --- | --- | --- |
| ***National ADI*** | | | | |
|  | β-value (95% CI)^c^ | *P*-value | β-value (95% CI)^c^ | *P*-value |
| 20% Most disadvantaged neighborhoods | 0.078 (-0.016 to 0.171) | .103 | -0.076 (-0.016 to 0.168) | .105 |
| Male sex | -0.047 (-0.085 to -0.009) | .016 | -0.052 (-0.090 to -0.015) | .006 |
| Age | -0.003 (-0.004 to -0.002) | < .001 | -0.003 (-0.004 to -0.002) | < .001 |
| WMHV | -0.005 (-0.007 to -0.002) | < .001 | -0.005 (-0.007 to -0.002) | < .001 |
| ***State ADI*** | | | | |
|  | β-value (95% CI)^c^ | *P*-value | β-value (95% CI)^c^ | *P*-value |
| 20% Most disadvantaged neighborhoods | 0.069 (-0.020 to 0.158) | .130 | 0.079 (-0.009 to 0.166) | .078 |
| Male sex | -0.047 (-0.085 to -0.009) | .015 | -0.053 (-0.090 to -0.015) | .005 |
| Age | -0.003 (-0.004 to -0.002) | < .001 | -0.003 (-0.004 to -0.002) | < .001 |
| WMHV | -0.005 (-0.007 to -0.002) | < .001 | -0.005 (-0.007 to -0.002) | < .001 |

Abbreviations: area disadvantage index (ADI), total white matter hyperintensity volume (WMHV).

^a^National ADI model: F(4,2822) = 14.22, *P* < .001, Adjusted R^2^ = 0.02; State ADI model: F(4,2822) = 14.13, *P* < .001, Adjusted R^2^ = 0.02.

^b^National ADI model: F(4,2822) = 20.59, *P* < .001, Adjusted R^2^ = 0.03; State ADI mode: F(4,2822) = 20.72, *P* < .001, Adjusted R^2^ = 0.03.

^c^Values presented are unstandardized.

**Table S8: Regression Model Results for Association Between Neighborhood-Level Disadvantage and Nucleus Accumbens Volume**

| **Variable** | **Left Hemisphere**^a^ | | **Right Hemisphere**^b^ | |
| --- | --- | --- | --- | --- |
| ***National ADI*** | | | | |
|  | β-value (95% CI)^c^ | *P*-value | β-value (95% CI)^c^ | *P*-value |
| 20% Most disadvantaged neighborhoods | -0.004 (-0.023 to -0.015) | .670 | -0.006 (-0.025 to 0.013) | .565 |
| Male sex | -0.020 (-0.028 to -0.013) | < .001 | -0.007 (-0.015 to 0.0003) | .060 |
| Age | -0.0009 (-0.001 to -0.0007) | < .001 | -0.001 (-0.001 to -0.0009) | < .001 |
| WMHV | -0.002 (-0.003 to -0.002) | < .001 | -0.002 (-0.003 to -0.002) | < .001 |
| ***State ADI*** | | | | |
|  | β-value (95% CI)^c^ | *P*-value | β-value (95% CI)^c^ | *P*-value |
| 20% Most disadvantaged neighborhoods | -0.008 (-0.026 to 0.009) | .354 | -0.006 (-0.024 to 0.012) | .506 |
| Male sex | -0.020 (-0.028 to -0.012) | < .001 | -0.007 (-0.015 to 0.0004) | .062 |
| Age | -0.0009 (-0.001 to -0.0007) | < .001 | -0.001 (-0.001 to -0.0009) | < .001 |
| WMHV | -0.002 (-0.003 to -0.002) | < .001 | -0.002 (-0.003 to -0.002) | < .001 |

Abbreviations: area disadvantage index (ADI), total white matter hyperintensity volume (WMHV).

^a^National ADI model: F(4,2822) = 51.71, *P* < .001, Adjusted R^2^ = 0.07; State ADI model: F(4,2822) = 51.89, *P* < .001, Adjusted R^2^ = 0.07.

^b^National ADI model: F(4,2822) = 54.15, *P* < .001, Adjusted R^2^ = 0.07; State hemisphere: F(4,2822) = 54.18, *P* < .001, Adjusted R^2^ = 0.07.

^c^Values presented are unstandardized.

**Table S9: Regression Model Results for Association Between Neighborhood-Level Disadvantage and Anterior Cingulate Cortex Volume**

| **Variable** | **Left Hemisphere**^a^ | | **Right Hemisphere**^b^ | |
| --- | --- | --- | --- | --- |
| ***National ADI*** | | | | |
|  | β-value (95% CI)^c^ | *P*-value | β-value (95% CI)^c^ | *P*-value |
| 20% Most disadvantaged neighborhoods | -0.059 (-0.196 to 0.077) | .393 | 0.019 (-0.126 to 0.164) | .796 |
| Male sex | 0.036 (-0.019 to 0.092) | .202 | 0.006 (-0.053 to 0.065) | .839 |
| Age | -0.002 (-0.004 to -0.0009) | .001 | -0.00003 (-0.002 to 0.002) | .972 |
| WMHV | -0.006 (-0.010 to -0.002) | .001 | -0.012 (-0.016 to -0.008) | < .001 |
| ***State ADI*** | | | | |
|  | β-value (95% CI)^c^ | *P*-value | β-value (95% CI)^c^ | *P*-value |
| 20% Most disadvantaged neighborhoods | -0.064 (-0.194 to 0.066) | .336 | 0.022 (-0.116 to 0.160) | .756 |
| Male sex | 0.037 (-0.019 to 0.092) | .197 | 0.006 (-0.053 to 0.065) | .843 |
| Age | -0.002 (-0.004 to -0.0009) | .001 | -0.00003 (-0.002 to 0.002) | .974 |
| WMHV | -0.006 (-0.010 to -0.002) | .001 | -0.012 (-0.012 to -0.008) | < .001 |

Abbreviations: area disadvantage index (ADI), total white matter hyperintensity volume (WMHV).

^a^National ADI model: F(4,2822) = 6.03, *P* < .001, Adjusted R^2^ = 0.007; State ADI model: F(4,2822) = 6.08, *P* < .001, Adjusted R^2^ = 0.007.

^b^National ADI model: F(4,2822) = 8.45, *P* < .001, Adjusted R^2^ = 0.01; State ADI model: F(4,2822) = 8.46, *P* < .001, Adjusted R^2^ = 0.01

^c^Values presented are unstandardized.

**Table S10: Regression Model Results for Association Between Neighborhood-Level Disadvantage and Posterior Cingulate Cortex Volume**

| **Variable** | **Left Hemisphere**^a^ | | **Right Hemisphere**^b^ | |
| --- | --- | --- | --- | --- |
| ***National ADI*** | | | | |
|  | β-value (95% CI)^c^ | *P*-value | β-value (95% CI)^c^ | *P*-value |
| 20% Most disadvantaged neighborhoods | 0.099 (-0.001 to -0.198) | .052 | 0.038 (-0.067 to 0.143) | .475 |
| Male sex | -0.077 (-0.117 to -0.036) | < .001 | -0.046 (-0.088 to -0.003) | .037 |
| Age | -0.006 (-0.007 to -0.005) | < .001 | -0.006 (-0.007 to -0.005) | < .001 |
| WMHV | -0.008 (-0.011 to -0.005) | < .001 | -0.004 (-0.007 to -0.002) | .002 |
| ***State ADI*** | | | | |
|  | β-value (95% CI)^c^ | *P*-value | β-value (95% CI)^c^ | *P*-value |
| 20% Most disadvantaged neighborhoods | 0.074 (-0.021 to 0.169) | .125 | 0.074 (-0.025 to 0.174) | .143 |
| Male sex | -0.077 (-0.117 to -0.036) | < .001 | -0.047 (-0.089 to -0.004) | .033 |
| Age | -0.006 (-0.007 to -0.005) | < .001 | -0.006 (-0.007 to -0.005) | < .001 |
| WMHV | -0.008 (-0.011 to -0.005) | < .001 | -0.004 (-0.007 to -0.002) | .002 |

Abbreviations: area disadvantage index (ADI), total white matter hyperintensity volume (WMHV).

^a^National ADI model: F(4,2822) = 45.21, *P* < .001, Adjusted R^2^ = 0.06; State ADI model: F(4,2822) = 44.84, *P* < .001, Adjusted R^2^ = 0.06.

^b^National ADI model: F(4,2822) = 31.06, *P* < .001, Adjusted R^2^ = 0.04; State ADI model: F(4,2822) = 31.49, *P* < .001, Adjusted R^2^ = 0.04.

^c^Values presented are unstandardized.

**Table S11: Regression Model Results for Association Between Neighborhood-Level Disadvantage and Medial Prefrontal Cortex Volume**

| **Variable** | **Left Hemisphere**^a^ | | **Right HemisphereI**^b^ | |
| --- | --- | --- | --- | --- |
| ***National ADI*** | | | | |
|  | β-value (95% CI)^c^ | *P*-value | β-value (95% CI)^c^ | *P*-value |
| 20% Most disadvantaged neighborhoods | -0.030 (-0.085 to 0.025) | .286 | 0.024 (-0.035 to 0.082) | .426 |
| Male sex | -0.016 (-0.038 to 0.006) | .158 | 0.007 (-0.017 to 0.031) | .574 |
| Age | -0.002 (-0.002 to -0.001) | < .001 | -0.002 (-0.003 to -0.001) | < .001 |
| WMHV | -0.004 (-0.005 to -0.002) | < .001 | -0.006 (-0.008 to -0.005) | < .001 |
| ***State ADI*** | | | | |
|  | β-value (95% CI)^c^ | *P*-value | β-value (95% CI)^c^ | *P*-value |
| 20% Most disadvantaged neighborhoods | -0.040 (-0.093 to 0.012) | .128 | 0.012 (-0.043 to 0.068) | .665 |
| Male sex | -0.016 (-0.038 to 0.007) | .169 | 0.007 (-0.017 to 0.031) | .569 |
| Age | -0.002 (-0.002 to -0.001) | < .001 | -0.002 (-0.003 to -0.001) | < .001 |
| WMHV | -0.004 (-0.005 to -0.002) | < .001 | -0.006 (-0.008 to -0.005) | < .001 |

Abbreviations: area disadvantage index (ADI), total white matter hyperintensity volume (WMHV).

^a^National ADI model: F(4,2822) = 16.29, *P* < .001, Adjusted R^2^ = 0.02; State ADI model: F(4,2822) = 16.59, *P* < .001, Adjusted R^2^ = 0.02.

^b^National ADI model: F(4,2822) = 28.29, *P* < .001, Adjusted R^2^ = 0.04; State ADI model: F(4,2822) = 28.17, *P* < .001, Adjusted R^2^ = 0.04.

^c^Values presented are unstandardized.

**Table S12: Regression Model Results for Association Between Neighborhood-Level Disadvantage and Lateral Prefrontal Cortex Volume**

| **Variable** | **Left Hemisphere**^a^ | | **Right Hemisphere**^b^ | |
| --- | --- | --- | --- | --- |
| ***National ADI*** | | | | |
|  | β-value (95% CI)^c^ | *P*-value | β-value (95% CI)^c^ | *P*-value |
| 20% Most disadvantaged neighborhoods | -0.351 (-1.008 to 0.305) | .294 | -0.606 (-1.252 to 0.039) | .065 |
| Male sex | -0.115 (-0.383 to 0.153) | .399 | 0.010 (-0.254 to 0.273) | .942 |
| Age | -0.037 (-0.044 to -0.030) | < .001 | -0.030 (-0.037 to -0.023) | < .001 |
| WMHV | -0.113 (-0.131 to -0.095) | < .001 | -0.096 (-0.114 to -0.079) | < .001 |
| ***State ADI*** | | | | |
|  | β-value (95% CI)^c^ | *P*-value | β-value (95% CI)^c^ | *P*-value |
| 20% Most disadvantaged neighborhoods | -0.435 (-1.060 to 0.189) | .172 | -0.584 (-1.198 to 0.030) | .062 |
| Male sex | -0.111 (-0.379 to 0.157) | .416 | 0.013 (-0.251 to 0.276) | .923 |
| Age | -0.037 (-0.044 to -0.030) | < .001 | -0.030 (-0.037 to -0.024) | < .001 |
| WMHV | -0.113 (-0.131 to -0.094) | < .001 | -0.096 (-0.114 to -0.079) | < .001 |

Abbreviations: area disadvantage index (ADI), total white matter hyperintensity volume (WMHV).

^a^National ADI model: F(4,2822) = 74.62, *P* < .001, Adjusted R^2^ = 0.09; State ADI model: F(4,2822) = 74.83, *P* < .001, Adjusted R^2^ = 0.09.

^b^National ADI model: F(4,2822) = 54.46, *P* < .001, Adjusted R^2^ = 0.07; State ADI model: F(4,2822) = 54.48, *P* < .001, Adjusted R^2^ = 0.07.

^c^Values presented are unstandardized.
